# Causal integration of multi-omics data with prior knowledge to generate mechanistic hypotheses

**DOI:** 10.1101/2020.04.23.057893

**Authors:** Aurelien Dugourd, Christoph Kuppe, Marco Sciacovelli, Enio Gjerga, Kristina B. Emdal, Dorte B. Bekker-Jensen, Jennifer Kranz, Eric. M. J. Bindels, Ana S. H. Costa, Jesper V. Olsen, Christian Frezza, Rafael Kramann, Julio Saez-Rodriguez

## Abstract

Multi-omics datasets can provide molecular insights beyond the sum of individual omics. Diverse tools have been recently developed to integrate such datasets, but there are limited strategies to systematically extract mechanistic hypotheses from them. Here, we present COSMOS (Causal Oriented Search of Multi-Omics Space), a method that integrates phosphoproteomics, transcriptomics, and metabolics datasets. COSMOS combines extensive prior knowledge of signaling, metabolic, and gene regulatory networks with computational methods to estimate activities of transcription factors and kinases as well as network-level causal reasoning. COSMOS provides mechanistic hypotheses for experimental observations across multi-omics datasets. We applied COSMOS to a dataset comprising transcriptomics, phosphoproteomics, and metabolomics data from healthy and cancerous tissue from nine renal cell carcinoma patients. We used COSMOS to generate novel hypotheses such as the impact of Androgen Receptor on nucleoside metabolism and the influence of the JAK-STAT pathway on propionyl coenzyme A production. We expect that our freely available method will be broadly useful to extract mechanistic insights from multi-omics studies.

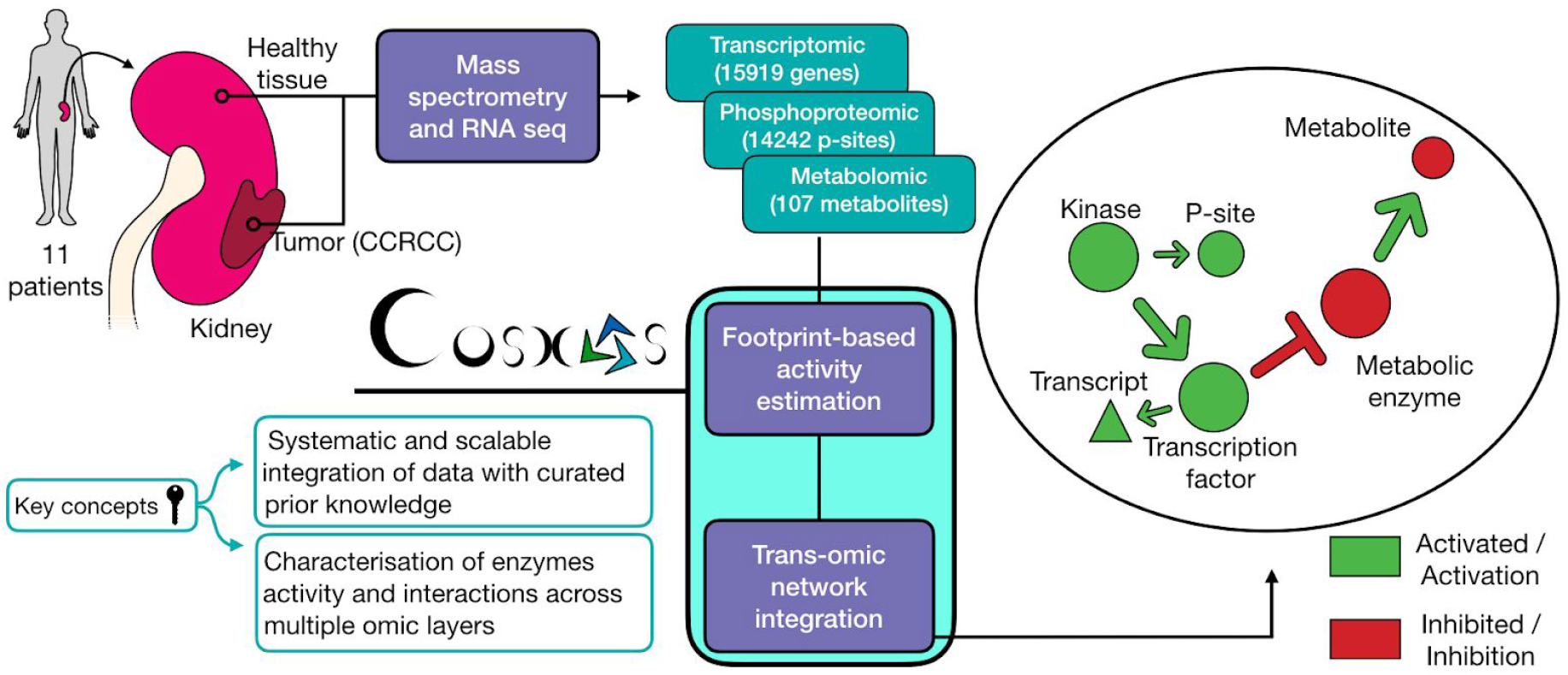

## 1. Introduction

“Omics” technologies measure at the same time thousands of biological molecules in biological samples, from DNA, RNA and proteins to metabolites. Omics datasets are an essential component of systems biology, and are made possible by the popularization of analytical methods such as Next Generation Sequencing or Mass-Spectrometry. Omics data have enabled the unbiased characterization of the molecular features of multiple human diseases, particularly in cancer^1–3^. It is becoming increasingly common to characterize multiple omics layers in parallel, with so-called “trans-omics analysis”, to gain biological insights spanning multiple types of cellular processes^4–6^. Consequently, many tools are developed to analyze such data^7–11^, mainly by adapting and combining existing “single omics” methodologies to multiple parallel datasets. These methods identify groups of measurements and derive integrated statistics to describe them, effectively reducing the dimensionality of the datasets. These methods are useful to provide a global view on the data, but additional processing is required to extract mechanistic insights from them.

To extract mechanistic insights from datasets, some methods (such as pathway enrichment analysis) use prior knowledge about the players of the process being investigated. For instance, differential changes in the expression of the genes that constitute a pathway gene expression are used to infer the activity of that pathway. Methods that a priori define groups of measurements based on known regulated targets (that we call footprints^12^) of transcription factors (TFs)^13,14^, kinases/phosphatases^15^ and pathway perturbations^16^, provide integrated statistics that can be interpreted as a proxy of the activity of a molecule or process. These methods seem to estimate more accurately the status of processes than classic pathway methods ^12,16,17^. Since each of these types of footprint methods work with a certain type of omics data, finding links between them could help to interpret them collectively in a mechanistic manner. For example, one can use a network diffusion algorithm, such as TieDIE^18^, to connect different omics footprints together^19^. This approach provides valuable insights, but diffusion (or random walk) based algorithms do not typically take into account causal information (such as activation/inhibition) that is available and are important to extract mechanistic information. TieDIE partially addressed this problem by focusing the diffusion process on causally coherent subparts of a network of interest, but it is thus limited to local causality.

Recently, we proposed the CARNIVAL tool^20^ to systematically generate mechanistic hypotheses connecting TFs through global causal reasoning supported by Integer Linear Programming. CARNIVAL connects activity perturbed nodes such as drug targets with deregulated TFs activities by contextualizing a signed and directed Prior Knowledge Network (PKN). We had hypothesized how such a method could potentially be used to actually connect footprint based activity estimates across multiple omics layers^12^.

In this study, we introduce COSMOS (Causal Oriented Search of Multi-Omics Space), an approach that builds on CARNIVAL to connect TF and kinase/phosphatases activities as well as metabolite abundances with a novel PKN spanning across multiple omics layers (Figure 1). COSMOS uses CARNIVAL’s Integer Linear Programming (ILP) optimization strategy to find the smallest coherent subnetwork causally connecting as many deregulated TFs, kinases/phosphatases and metabolites as possible. The subnetwork is extracted from a novel integrated PKN spanning signaling, transcriptional regulation and metabolism of > 67000 edges. CARNIVAL’s ILP formulation effectively allows to evaluate the entire network’s causal coherence given a set of known TF, kinases/phosphatases activities and metabolite abundances. While we showcase this method using transcriptomics, phosphoproteomics and metabolomics inputs, COSMOS can theoretically be used with any other additional inputs, as long as they can be linked to functional insights (for example, a set of deleterious mutations). As a case study, we generated transcriptomics, phosphoproteomics, and metabolomics datasets from kidney tumor tissue and corresponding healthy kidney tissue out of nine clear cell renal cell carcinoma (ccRCC) patients. We estimated changes of activities of TFs and kinase/phosphatases as well as metabolite abundance differences between tumor and healthy tissue. We integrated multiple curated resources of interactions between proteins, transcripts and metabolites together to build a trans-omics PKN. Next, we contextualized the trans-omics PKN to a specific experiment. To do so, we identified causal pathways from our prior knowledge that connect the observed changes in activities of TFs, kinases, phosphatases and metabolite abundances between tumor and healthy tissue. These causal pathways can be used as hypothesis generation tools to better understand the molecular phenotype of kidney cancer.

**Figure 1.**
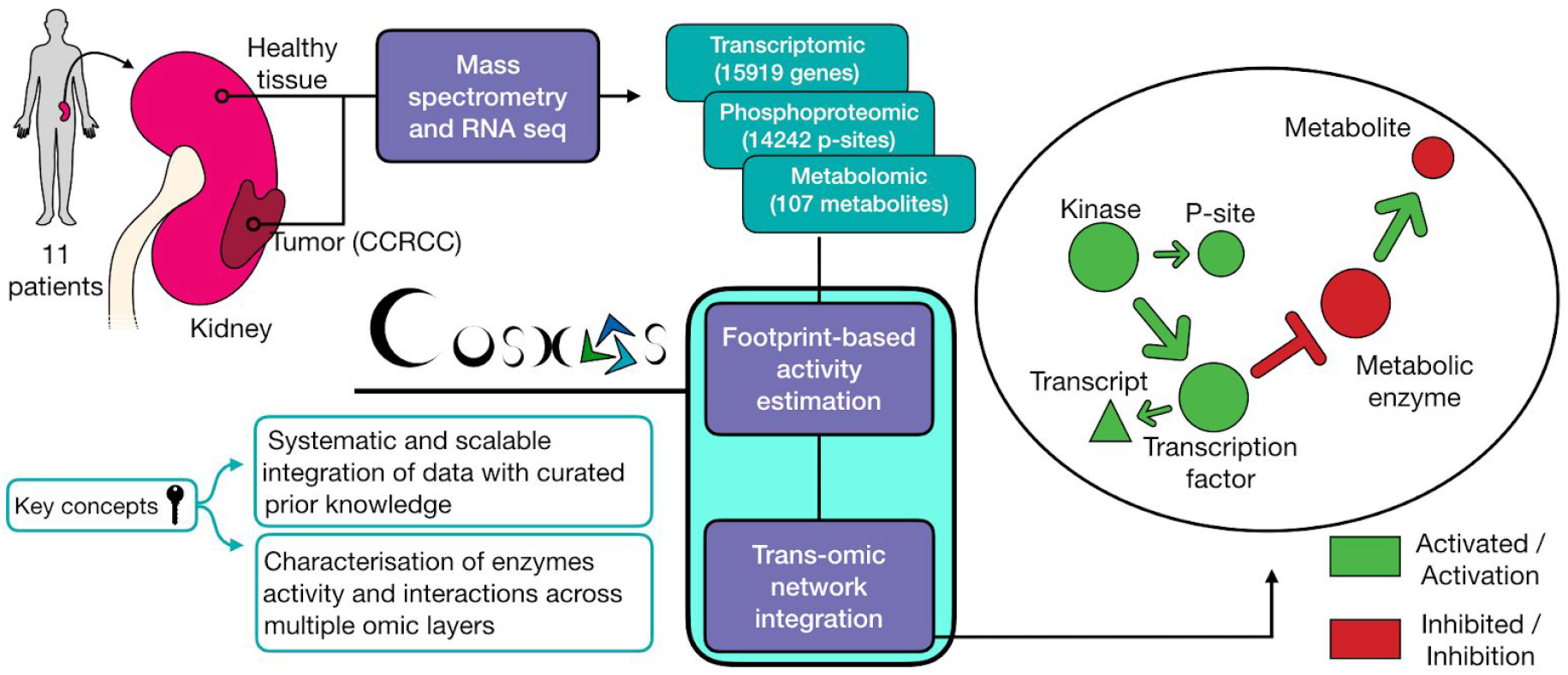
Overview of analysis pipeline. From left to right: We sampled and processed 11 patient tumors and healthy kidney tissues from the same kidney through RNA sequencing and 9 of those same patients through mass-spectrometry to characterise their transcriptomics, phospho-proteomics, and metabolomics profiles. We calculated differential abundance for each detected gene, phospho-peptide and metabolite. We estimated kinase and transcription factor activities using the differential analysis statistics and footprint-based methods. We used the estimated activities alongside the differential metabolite abundances to contextualise (i.e. extract the subnetwork that better explains the phenotype of interest) a generic trans-omics causal network.

## 2. Results

### 2.1 Building the multi-omics dataset

To build a multi-omics dataset of renal cancer, we performed transcriptomics, phosphoproteomics, and metabolomics analyses of renal nephrectomies and adjacent normal tissues of renal cancer patients (for details on the patients see methods). First, we processed the different omics datasets to prepare for the analysis. For the transcriptomics dataset, 15919 transcripts with average counts > 50 were kept for subsequent analysis. In the phosphoproteomics dataset, 14243 phosphosites detected in at least four samples were kept. In the metabolomics dataset 107 metabolomics detected across 16 samples were kept. Principal Component Analysis (PCA) of each omics dataset independently showed a clear separation of healthy and tumor tissues on the first component (transcriptomics : 40% of explained variance (EV), phosphoproteomics : 26% of EV, metabolomics : 28% of EV, Supplementary Figure 1), suggesting that tumor sample displayed molecular deregulations spanning across signaling, transcription and metabolism. Each omics dataset was independently submitted to differential (tumor vs healthy tissue) analysis using LIMMA^21^. We obtained 6699 transcript and 21 metabolites significantly regulated with False Discovery Rate (FDR) < 0.05. While only 11 phosphosites were found under 0.05 FDR, 447 phosphosites had an FDR < 0.2. This result confirmed that tumor samples displayed molecular deregulations spanning across signaling, transcription, and metabolism but that TF dysregulation is more pervasive. The differential statistics for all transcripts, phospho-proteins and metabolites were then used for further downstream analysis.

### 2.3 Footprint based transcription factor, kinase and phosphatase activity estimation

We then performed computational footprint analysis to estimate the activity of proteins responsible for changes observed in specific omics datasets. For transcriptomics and phosphoproteomics data, this analysis estimates transcription factor and kinases/phosphatase activity, respectively. 32586 Transcription Factor (TF) to target interactions (i. e. transcript under the direct regulation of a transcription factor) were obtained from DOROTHEA^14^, a meta-resource of TF-target interactions. Those TF-target interactions span over 452 unique transcription factors. In parallel, 33616 interactions of kinase/phosphosphate and their phosphosite targets (i. e. phosphopeptides directly (de)phosphorylated by specific kinases(phosphatases)) were obtained from Omnipath^22^ kinase substrate network, a meta resource focused on curated information on signaling processes. Only TFs and kinases/phosphatases with at least 25 and 5 detected substrates, respectively, were included. This led to the activity estimation of 229 TFs and 174 kinases. In line with the results of the differential analysis, where fewer phosphosites were deregulated than transcripts, TF activities displayed a stronger deregulation than kinases. TF activity scores reached a maximum of eight standard deviations (sd) for Transcription Factor AP-2 Gamma (TFAP2C) (compared to the null score distribution) while kinase activity scores reached a maximum of 4.6 sd for Casein Kinase 2 Alpha 1 (CSNK2A1). In total, 102 TFs and kinases/phosphatase had an absolute score over 1.7 sd (p-val<0.05) and were considered significantly deregulated in kidney tumor samples. The presence of several known signatures of ccRCC corroborated the validity of our analysis. For instance, hypoxia (HIF1A, EPAS1), inflammation (STAT1/2) and oncogenic (MYC, Cyclin Dependent Kinase 2 and 7 (CDK2/7)) markers were up-regulated in tumors compared to healthy tissues (Figure 2). Furthermore, among suppressed TFs we identified, HNF4A has been previously associated with ccRCC^23^.

**Figure 2.**
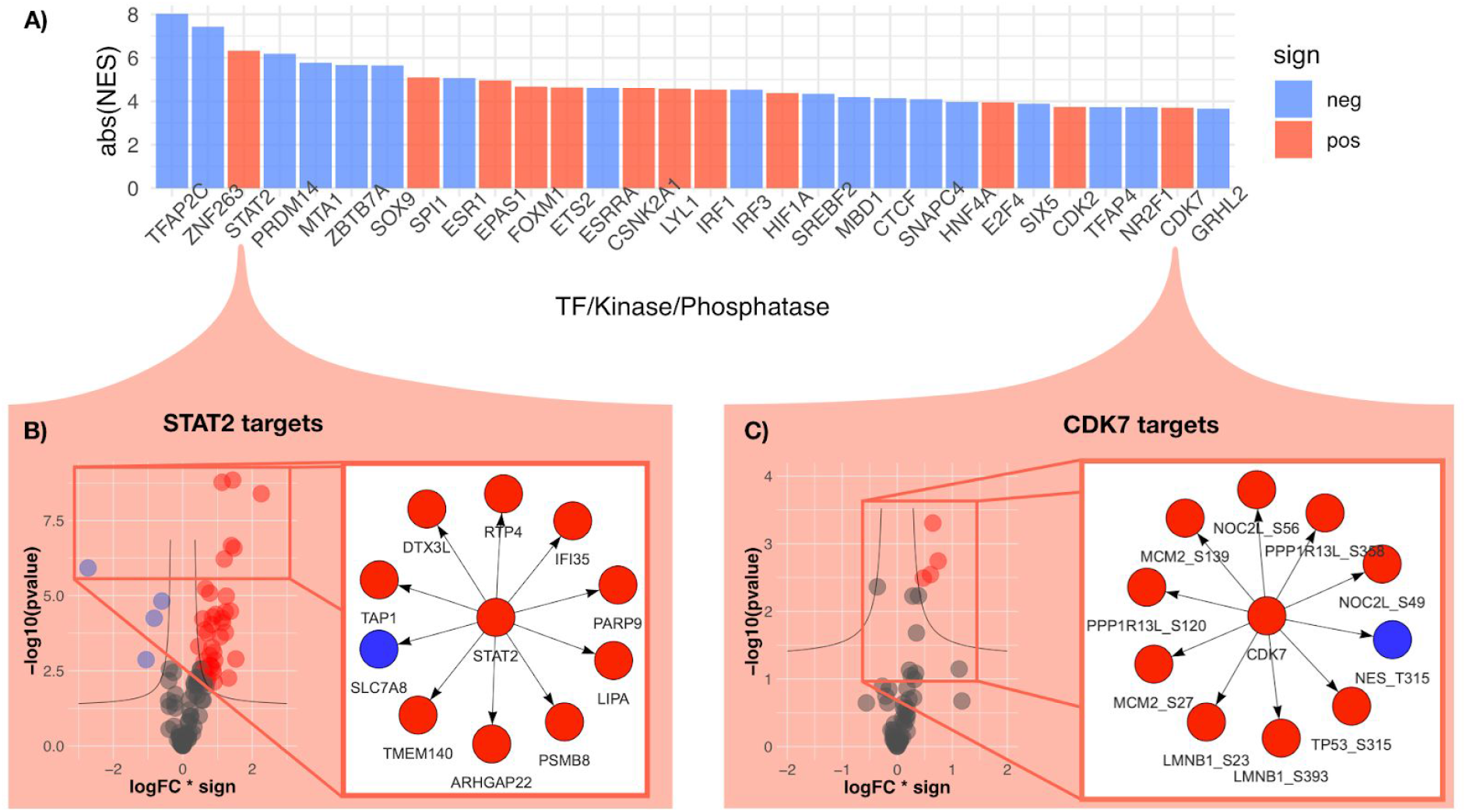
TF, kinase and phosphatase activities that change the most between cancer and healthy tissue. A) Bar plot displaying the Normalised Enrichment Score (NES, proxy of activity change) of the 30 most changing TF, kinase and phosphatases activities between kidney tumor and adjacent healthy tissue. Blue/red color represent the sign of the activity change (negative/positive, respectively). B) Right panel shows the 10 most changing RNA abundances of the STAT2 regulated transcripts. Left panel shows the change of abundances of all STAT2 regulated transcripts that were used to estimate its activity change. X axis represents log fold change of regulated transcripts multiplied by the sign of regulation (−1 for inhibition and 1 for activation of transcription). Y axis represents the significance of the log fold change (−log10 of p-value). C) Right panel shows the 10 most changing phospho-peptide abundances of the CDK7 regulated phospho-peptides. Left panel shows the change of abundances of all CDK7 regulated phospho-peptides that were used to estimate its activity change.

### 2.4 Causal network analysis

We set out to find potential causal mechanistic pathways that could explain the changes we observed in TF, kinases/phosphatase activities, and metabolic abundances. Thus, we developed a systematic approach to search in public databases, via OmniPath, for plausible causal links between significantly deregulated TFs, kinases/phosphatases and metabolites. In brief, we investigated if changes in TF, kinase/phosphatase activities, and metabolite abundance can explain each other with the support of literature-curated molecular interactions. An example of such a mechanism can be the activation of the transcription of MYC gene by STAT1. Since both STAT1 and MYC display increased activities in tumors, and there is evidence in the literature that STAT1 can regulate MYC transcription^24,25^, it may indicate that this mechanism is responsible for this observation.

First, we needed to map the deregulated TFs, kinases and metabolites on a causal prior knowledge network spanning over signaling pathways, gene regulation, and metabolic networks. Hence, we combined multiple sources of experimentally curated causal links together to build a trans-omics causal prior knowledge network (trans-omics PKN). This trans-omics PKN must include direct causal links between proteins (kinase to kinase, TF to kinase, TF to metabolic enzymes, etc…), between proteins and metabolites (reactants to metabolic enzymes and metabolic enzymes to products) and between metabolites and proteins (allosteric regulations). High confidence (>= 900 combined score) allosteric regulations of the STITCH database^26^ were used as the source of causal links between metabolites and enzymes (Figure 3A). The directed signed interactions of the Omnipath database were used as a source of causal links between proteins (Figure 3B). The human metabolic network Recon3D^27^ (without cofactors and hyper-promiscuous metabolites, see methods) was converted to a causal network and used as the source of causal links between metabolites and metabolic enzymes (Figure 3C). The resulting trans-omics PKN consists of 69517 interactions and contains causal paths linking TF/kinase/phosphatase with metabolites and vice-versa in a machine readable format. This network is available at http://metapkn.omnipathdb.org/.

**Figure 3.**
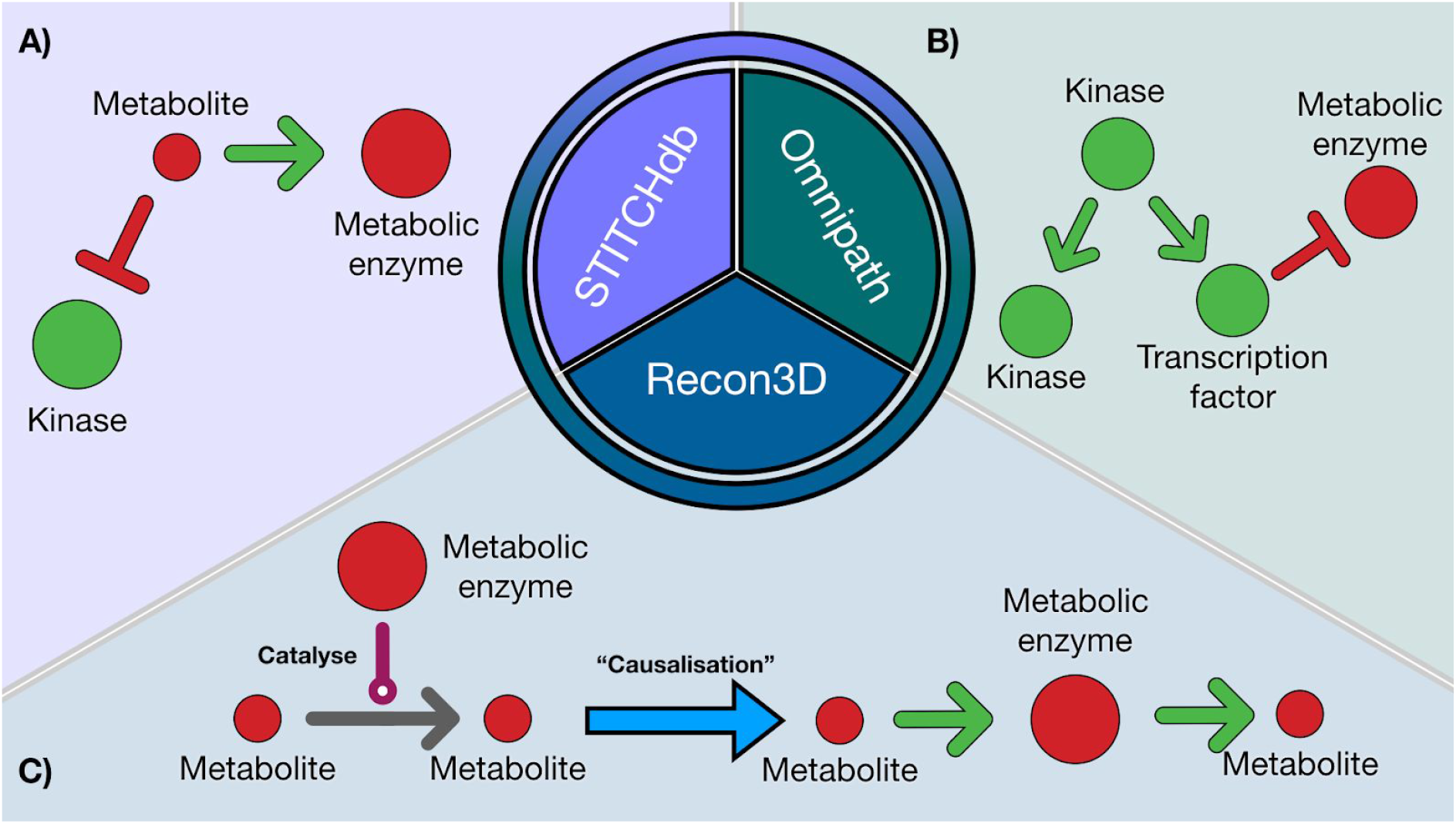
Graphical explanation of trans-omics PKN sources. Schematic representation of the trans-omics generic network (trans-omics PKN) created combining STITCHdb, Omnipath and Recon3D. A) STITCHdb provides information on inhibition/activation of enzyme activities mediated by metabolites. B) Omnipath provides information inhibition/activation of enzyme activities mediated by other enzymes based mainly on curated resources. C) Recon3D provides information on reactants and products associated with metabolic enzymes. To make this information compatible with the causal edges from Omnipath and STITCH, the interactions of recon3D are converted so that reactants “activate” their metabolic enzymes, which themselves “activate” their products.

We then used the trans-omics PKN to systematically search causal paths between the deregulated TFs, kinases/phosphatases and metabolites. The CARNIVAL^20^ tool uses Integer Linear Programming (ILP) to find causal paths between perturbations and deregulated TFs using a PKN and infers the state of intermediate nodes when it is unknown. Here we use CARNIVAL with our trans-omics PKN to find the smallest sign-coherent subnetwork connecting as many deregulated TFs, kinases/phosphatases, and metabolites as possible. CARNIVAL is first used to find causal paths going from TFs/kinases/phosphatases to the metabolites (the ‘forward network’). Then, in order to complete the loop, CARNIVAL is used to go from metabolites to TFs/Kinases/phosphatases (‘backward network’).

When applied to our kidney cancer data, the two resulting (forward and backward) networks are then combined into a single network of 250 signed directed interactions (Supplementary Figure 2). These interactions are directly interpretable as mechanistic hypotheses. We present some of them using official symbol nomenclature for genes and metabolites. For example, it appears that Androgen Receptor (AR) activity inhibition could be responsible for the observed downregulation of uridine, adenine, and inosine metabolism by down-regulating the expression of ACPP, DBI, and SMS metabolic enzymes (Figure 4A). Of note, AR expression has a protective role in ccRCC progression ^28,29^. Interestingly, the COSMOS network shows adenine depletion could lead to adenosine depletion (since adenosine can be produced from adenine). Adenosine is a known activator of the C-X-C Motif Chemokine Receptor 4 (CXCR4) ^30,31^, so its depletion could lead to the predicted down-regulation of CXCR4 activity. The combined AR and CXCR4 down-regulation might indicate that these tumors are not metastatic ^32–34^. The COSMOS network also shows that CXCR4 regulates Phosphatidylinositol-4,5-Bisphosphate 3-Kinase Catalytic Subunit Gamma (PIK3CG), which itself regulates 3-Phosphoinositide Dependent Protein Kinase 1 (PDPK1). Thus, CXCR4 down-regulation could then explain PDPK1 activity down-regulation through the inhibition of PIK3CG. COSMOS further proposes that the activation of JAK kinase would be a good explanation for the apparent activation of STAT transcription factors in the tumor, leading to activation of IRF1 and MYC (Figure 4B). Interestingly, JAK2 was found to be amplified in ccRCC ^35^. The STAT3 activation could explain the depletion of o-propanoylcarnitine due to the downregulation of metabolic enzymes responsible for the transport of its precursor, Sterol Carrier Protein 2 (SCP2). CDK2 could itself explain the activity of ATM and TP53 through Forkhead Box M1 (FOXM1) and Aurora Kinase B (AURKB) signaling, leading to the activation of Dual specificity tyrosine-phosphorylation-regulated kinase 2 (DYRK2), HIF1A and the accumulation of L-Glutamine (Figure 4C). FOXM1 was recently highlighted as a particularly important driver of metabolic changes in ccRCC^36^. Finally, the COSMOS network shows that the down-regulation of PDPK1 appears as a good explanation for L-Citrulline accumulation and ethanolamine depletion, by indirectly modulating the activity of Nitric Oxide Synthase 1 (NOS1) and Phospholipase D1 (PLD1) metabolic enzymes (Figure 4D). Furthermore, PDPK1 directly controls the activity of the Protein Kinase C protein family (PRKCA, PRKCD, PRKCE and PRKACA). These kinases are known to be involved in metastasis progression ^37,38^. Their down-regulation predicted by COSMOS further supports the idea that these tumors are not metastatic. These results demonstrate how the pipeline can be used to extract relevant mechanistic hypotheses explaining the enzymatic and metabolic deregulations at signaling and transcriptional levels.

**Figure 4.**
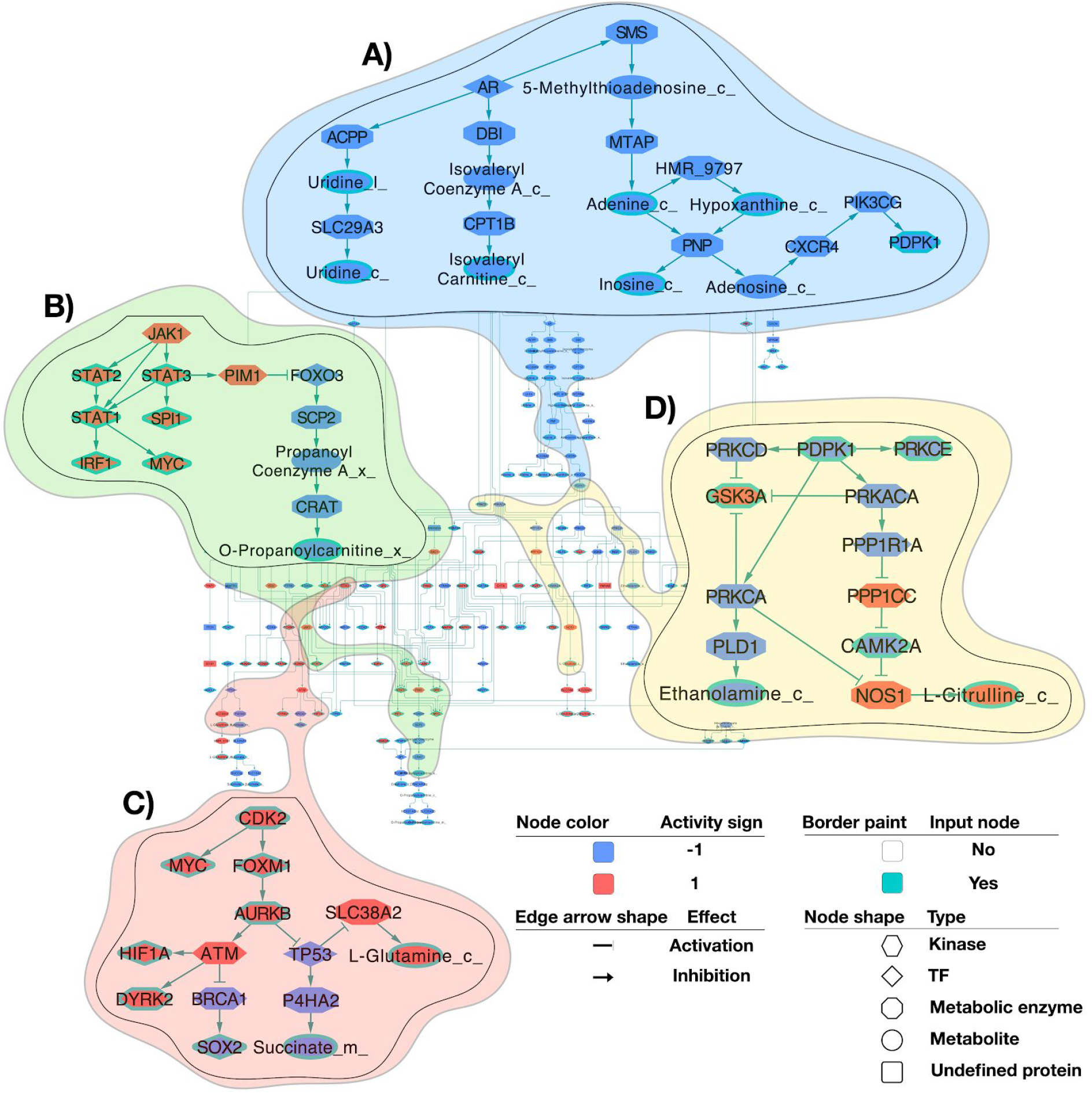
Systematically generated mechanistic hypotheses explaining changing TF, kinase, phosphatase activities and metabolic abundances. COSMOS generates mechanistic hypotheses which are represented in the form of a context specific causal network. This network links the significant changes in estimated enzyme activities and metabolic abundance (192 nodes and 250 edges). Diamond shapes represent TFs, hexagon shape represents kinases and phosphatases, octagon shape represents metabolic enzymes and ellipse shape represents metabolites. Blue/red color represents inhibited/activated enzyme activities and depleted/accumulated metabolites. Edges with arrowheads represent activatory interactions and T-shaped ones represent inhibitory interactions. A full-page version of this figure is available as Supplementary Figure 2 A), B), C) and D) represent subnetworks extracted to zoom on specific hypotheses. For example, B) represents how AR activity can lead to the inhibition of nucleotides and isovaleryl carnitine synthesis observed in tumors compared to healthy tissues, in turn explaining the inhibited activity of PDPK1.

### 2.5 Consistency analysis

Due to the combined effect of experimental noise and incompleteness of prior knowledge (kinase/substrate interactions, TF/targets interactions and meta PKN), it is critical to assess the performance of the pipeline presented above. We first looked if some of the generated hypotheses (see 2.3) were supported by parts of the datasets that were not directly used by CARNIVAL (Supplementary Figure 3). We couldn’t estimate the True Negative Rate of CARNIVAL in this multi-omics context. Indeed, nodes that are not integrated in the final subnetwork by CARNIVAL are simply not considered informative to explain the relationship between the input protein activities and metabolites. Yet, that doesn’t inform us on their actual functional state. Consequently, we focused on the True Positive Rate (TPR), for which we had reasonable estimates. The TF activity displayed by CARNIVAL can come from two distinct sources. The first source consists of the original footprint based activity estimation (using DOROTHEA and transcript abundances of target genes). The second source consists of actual CARNIVAL activity predictions based on molecular signal propagation through activating/inhibiting links (not using transcript abundances) connecting TF, kinases and phosphatases together. This is the case as some TFs can serve as intermediate links to connect upstream perturbations with downstream nodes. Consequently, in the case of a TF, CARNIVAL will implicitly model its action on the direct downstream targets. Thus, for every TF/target regulation of the CARNIVAL network, we checked whether the change of abundance of the target transcripts was actually coherent with the predicted activity of the TF displayed by CARNIVAL. We tested this over a range of differential transcript abundance t-value threshold between 0 and 2. Nine transcripts were regulated by TF whose activity was predicted by CARNIVAL only (second source), that is, the transcripts were not used as inputs to build the COSMOS model. Out of those nine transcripts, the TPR ranged between 0.62 and 0.15 depending on the t-value threshold (n = 13) (Supplementary Figure 4). It performed better than a random baseline for considered t-value thresholds ranging from 0 to 1.7.

Another way to estimate the performance is to check if the CARNIVAL mechanistic hypotheses correspond to correlations observed in tumor tissues. Thus, on the one hand, a topological driven coregulation network was generated from the CARNIVAL network. The assumption behind this network is that direct downstream targets of the same enzymes should be co-regulated. On the other hand, a data driven correlation network of TFs, kinases and phosphatases was generated from tumor tissues alone. Assuming thresholds of absolute values of correlation ranging between 0 and 1 to define true positive co-regulations, the comparison between the topological driven coregulation network and the data driven correlation network yielded a TPR of ranging between 0.6 and 0 (n = 157) for the carnival predictions (Supplementary Figure 4). It performed consistently better than a random baseline over the considered range of correlation coefficient thresholds. These results indicate that while some of the causal links predicted by CARNIVAL are potentially valid, some of them don’t find direct support in the data at hand. Thus, we sought to investigate if some of the mechanistic hypotheses could be experimentally validated.

## 3. Discussion

In this paper, we present COSMOS, an analysis pipeline to systematically generate mechanistic hypotheses by integrating multi-omics datasets with a broad range of curated resources of interactions between protein, transcripts and metabolites. We have first shown how TF, kinase and phosphatase activities could be coherently estimated from transcriptomics and phosphoproteomics datasets using footprint based analysis. This is a critical step before further mechanistic exploration. Indeed, transcript and phosphosite usually offer limited functional insights by themselves as their relationship with corresponding protein activity is usually not well characterised. Yet, they can provide information on the activity of the upstream proteins regulating their abundances. Thus, the functional state of kinases, phosphatases, and TFs is estimated from the observed abundance change of their known targets, i. e. their molecular footprint. Thanks to this approach, we could simultaneously characterise protein functional states in tumors at the level of signaling pathway and transcriptional regulation. Key actors of hypoxia response, inflammation pathway and oncogenic genes were found to have especially strong alteration of their functional states, such as HIF1A, EPAS1, STAT1/2, MYC and CDK2. Loss of VHL is a hallmark of ccRCC, and is directly linked to the stability of the HIF (HIF1A and EPAS1) proteins found deregulated by our analysis^39–41^. Finding these established signatures of ccRCC to be deregulated in our analysis is a confirmation of the validity of this approach.

We then used CARNIVAL with a novel trans-omics causal Prior Knowledge Network spanning signaling, transcription and metabolism to systematically find potential mechanisms linking deregulated protein activities and metabolite concentrations. To the best of our knowledge, this is the first attempt to integrate these three omics layers together in a systematic manner using causal reasoning. Previous methods studying signaling pathways with multi-omics quantitative datasets ^19^ connected TFs with kinases and they were limited by the preselected locally coherent subnetwork of the TieDIE algorithm. Introducing global causality with CARNIVAL along with metabolomics data allows us to obtain a direct mechanistic interpretation of links between proteins at different regulatory levels and metabolites. The goal of our approach is to find a coherent set of such mechanisms connecting as many of the observed deregulated protein activities and metabolite concentrations as possible. Using CARNIVAL is particularly interesting as all the proposed mechanisms between pairs of molecules (proteins and metabolites) have to be plausible not only in the context of their own pairwise interaction but also with respect to all other molecules that we wish to include in the model. For example, the proposed activation of MYC by STAT1 is further supported by IRF1 activation, because STAT1 is also known to activate IRF1. CARNIVAL allows us to scale this type of reasoning up to the entire PKN with all significantly deregulated protein activities and metabolites.

With our dataset, the resulting network showed that AR inhibition, a known tumor suppressor in kidney cancer ^42^, would be a good candidate to explain the inhibition of nucleotide metabolism. It also predicted a depletion of adenine and consequently the down-regulation of PDPK1 activity through CXCR4 (Figure 4A). Footprint analysis showed a down-regulation of PDPK1 (that is, the abundance of phosphorylation on its direct target phosphosites is decreasing) activity, which is surprising since its expression is usually associated with slower proliferation of kidney tumor cells^43,44^. Yet, the observed coordinated depletion of adenine, hypoxanthine and inosine strongly support the estimated down-regulation of PDPK1 activity. A consequence of PDPK1 activity down-regulation could also be the up-regulation of citrulline production by NOS1 (Figure 4C). COSMOS additionally predicted how JAK-STAT pathway activation could lead to an inhibition of the production of propanoyl-carnitine (Figure 4B). Diminution of carnitine and its derivative have been indeed previously observed in kidney cancer as a consequence of cachexia^45^. Finally we could show the importance of CDK2 as a master regulator of many kinases and transcription factors such as MYC, AURKB, E2F4 and consequently TP53 and ATM activities (Figure 4D). In particular, AURKB, which directly controls TP53 and ATM activities, appears to be a promising marker of kidney cancer^46–48^.

Then we assessed the performances of the approach in two ways. First, we used some of the data that was not directly used by CARNIVAL (i. e. genes that were not used for TF activity estimation and correlation between TF/kinase/phosphatase activities) to check the coherence of CARNIVAL predictions. Second, we used a tumor specific correlation network of TF and kinase activities to compare it to the co-regulation predicted by CARNIVAL. This yielded encouraging results, though imperfect, underscoring the fact that the mechanisms proposed by COSMOS - like those by any similar tool - are hypotheses.

There are three main known limits to the predictions of COSMOS. First, the input data is incomplete. Only a limited fraction of all potential phosphosites and metabolites are detected by mass spectrometry. This means that we have no information on a significant part of the PKN; part of the unmeasured network is kept in the analyses and the values are estimated as intermediate ‘hidden values’. Second, not all regulatory events between TFs, kinase and phosphatases and their targets are known, and activity estimation is based only on the known regulatory relationships. Thus, many TFs, kinase and phosphatases are not included because they have no curated regulatory interactions or no detected substrates in the data. Third, and conversely, COSMOS will find putative explanations within the existing prior knowledge that may not be the true mechanism, in particular if the latter is not captured in our knowledge.

These problems mainly originate from the importance that is given to prior knowledge in this method. Since prior knowledge is never perfect, the next steps of improvement could consist in finding ways to extract more knowledge from the observed data to weight in the contribution of prior knowledge. For instance, one could use the correlations between transcripts, phosphosites and metabolites to quantify the interactions available in databases such as Omnipath. Importantly, any other omics that relate to active molecules (such as miRNAs or metabolic enzyme fluxes) or can be used to estimate protein activities through footprint approaches (such as DNA accessibility or PTMs other than phosphorylation) can be seamlessly integrated. Moreover, COSMOS was designed to work with bulk omics datasets, and it will be very exciting to find ways of applying this approach to single cell datasets. Encouragingly, the footprint methods that bring data into COSMOS seem fairly robust to the characteristics of single-cell RNA data such as dropouts^49^. Finally, we expect that in the future data generation technologies will increase coverage and our prior knowledge will become more complete, reducing the mentioned limitations. In the meantime, we believe that COSMOS is already a useful tool to extract causal mechanistic insights from multi-omics studies.

## 4. Methods

### 4.1. Sample collection and processing

We included a total of 22 samples from 11 renal cancer patients (6 men, age 65.0+/−14.31, 5 women, age 65.2+/−9.257(mean+/−SD)) for transcriptomics and a subset of 18 samples from 9 of these patients (6 men, age 65+/−14.31; 3 women, age 63.33+/−11.06(mean+/−SD)) for metabolomics and phosphoproteomics analysis. Patients underwent nephrectomy due to renal cancer. We processed tissue from within the cancer and a distant unaffected area of the same kidney. The tissue was snap-frozen immediately after nephrectomy within the operation room. The clinical data of the included patients is outlined in supplementary table 1. Histological evaluation showed clear renal cell carcinoma in all patients.

#### Ethics

The local ethics committee of the University Hospital RWTH Aachen approved all human tissue protocols for this study (EK-016/17). The study was performed according to the declaration of Helsinki. Kidney tissues were collected from the Urology Department of the University Hospital Eschweiler from patients undergoing partial/- or nephrectomy due to kidney cancer. All patients gave informed consent.

#### Human tissue Processing

Kidney tissues were sampled by the surgeon from normal and tumor regions. The tissue was snap-frozen on dry-ice or placed in prechilled University of Wisconsin solution (#BTLBUW, Bridge to Life Ltd., Columbia, U.S.) and transported to our laboratory on ice.

#### RNA Isolation, library preparation, NGS sequencing

RNA was extracted according to the manufacturer ́s instructions using the RNeasy Mini Kit (QIAGEN). For rRNA-depleted RNA-seq using 1 and 10 ng of diluted total RNA, sequencing libraries were prepared with KAPA RNA HyperPrep Kit with RiboErase (Kapa Biosystems) according to the manufacturer’s protocol. Sequencing libraries were quantified using quantitative PCR (New England Biolabs, Ipswich, USA), equimolar pooled, final pool is normalized to 1,4 nM and denatured using 0.2 N NaOH and neutralized with 400nM Tris pH 8.0 prior to sequencing. Final sequencing was performed on a Novaseq6000 platform (IIlumina) according to the manufacturer’s protocols (Illumina, CA, USA).

#### Metabolomics

Snap-frozen tissue specimens were cut and weighed into Precellys tubes prefilled with ceramic beads (Stretton Scientific Ltd., Derbyshire, UK). An exact volume of extraction solution (30% acetonitrile, 50% methanol and 20% water) was added to obtain 40 mg specimen per mL of extraction solution. Tissue samples were lysed using a Precellys 24 homogeniser (Stretton Scientific Ltd., Derbyshire, UK). The suspension was mixed and incubated for 15 minutes at 4 °C in a Thermomixer (Eppendorf, Germany), followed by centrifugation (16,000 g, 15 min at 4°C). The supernatant was collected and transferred into autosampler glass vials, which were stored at −80 °C until further analysis.

Samples were randomised in order to avoid bias due to machine drift and processed blindly. LC-MS analysis was performed using a Q Exactive mass spectrometer coupled to a Dionex U3000 UHPLC system (both Thermo Fisher Scientific). The liquid chromatography system was fitted with a Sequant ZIC-pHILIC column (150 mm × 2.1 mm) and guard column (20 mm × 2.1 mm) from Merck Millipore (Germany) and temperature maintained at 45 °C. The mobile phase was composed of 20 mM ammonium carbonate and 0.1% ammonium hydroxide in water (solvent A), and acetonitrile (solvent B). The flow rate was set at 200 μL/min with the gradient described previously^50^. The mass spectrometer was operated in full MS and polarity switching mode. The acquired spectra were analysed using XCalibur Qual Browser and XCalibur Quan Browser software (Thermo Scientific).

#### Phosphoproteomics

Snap frozen tissues were heat inactivated (Denator T1 Heat Stabilizor, Denator) and transferred to a GndCl solution (6 M GndCl, 25 mM Tris, pH 8.5, Roche Complete Protease Inhibitor tablets (Roche)) and homogenized by ceramic beads using 2 steps of 20 s at 5500 rpm (Precellys 24, Bertin Technologies). The tissues were heated for 10 min at 95°C followed by micro tip sonication on ice and clarified by centrifugation (20 min, 16,000g, 4°C). Samples were reduced and alkylated by adding 5 mM tris(2-carboxyethyl)phosphine and 10 mM chloroacetamide for 20 min at room temperature.

Lysates were digested by Lys-C (Wako) in an enzyme/protein ratio of 1:100 (w/w) for 1 hour, followed by a dilution with 25 mM tris buffer (pH 8.5), to 2 M guanidine-HCl and further digested overnight with trypsin (Sigma-Aldrich; 1:100, w/w). Protease activity was quenched by acidification with TFA, and the resulting peptide mixture was concentrated on C18 Sep Pak Cartridges (Waters). Peptides were eluted with 40% ACN followed by 60% ACN. The combined eluate was reduced by SpeedVac, and the final peptide concentration was estimated by measuring absorbance at *A*280 on a NanoDrop (Thermo Fisher Scientific). Peptide (300 μg) from each sample was labeled with 1 of 11 different TMT reagents according to the manufacturer’s protocol (Thermo Fisher Scientific) for a total of four TMT sets. Each set comprised 10 samples and a common internal reference (composed of equal amounts of digested material from all samples).

After labeling, the samples were mixed and phosphopeptides were further enriched using titanium dioxide beads (5 μm Titansphere, GL Sciences, Japan). TiO2 beads were pre-incubated in 2,5-dihydroxybenzoic acid (20 mg/mL) in 80% ACN and 1% TFA (5 μL/mg of beads) for 20 min. Samples were brought to 80% ACN and 5% TFA. 1.5 mg beads (in 5μL of DHB solution) were added to each sample, which was then incubated for 20 min while rotating. After incubation, the beads were pelleted and fresh TiO2 beads were added to the supernatant for a second enrichment step. Beads were washed with five different buffers: (1) 80% ACN and 6% TFA, (2) 10% ACN and 6% TFA, (3) 80% ACN and 1% TFA, (4) 50% ACN and 1% TFA, (5) 10% ACN and 1% TFA. The final washing step was performed on a C8 stage tip, from which the phosphopeptides were with 20 μL 5% NH4OH followed by 20 μL 10% NH4OH with 25% ACN. Eluted peptides were fractionated using a reversed-phase Acquity CSH C18 1.7 μm 1 × 150 mm column (Waters, Milford, MA) on an UltiMate 3000 high-pressure liquid chromatography (HPLC) system (Dionex, Sunnyvale, CA) operating at 30 μL/min. Buffer A (5 mM ammonium bicarbonate) and buffer B (100% ACN) were used. Peptides were separated by a linear gradient from 5% B to 35% B in 55 min, followed by a linear increase to 70% B in 8 min and 12 fractions were collected in a concatenated manner.

The peptide solution was adjusted in volume to an appropriate concentration and kept in loading buffer (5% ACN and 0.1% TFA) prior to autosampling. An in-house packed 15 cm, 75 μm ID capillary column with 1.9 μm Reprosil-Pur C18 beads (Dr. Maisch, Ammerbuch, Germany) was used with an EASY-nLC 1200 system (Thermo Fisher Scientific, San Jose, CA). The column temperature was maintained at 40 °C using an integrated column oven (PRSO-V1, Sonation, Biberach, Germany) and interfaced online with a Q Exactive HF-X mass spectrometer. Formic acid (FA) 0.1% was used to buffer the pH in the two running buffers used. The gradients went from 8 to 24% acetonitrile (ACN) in 50 minutes, followed by 24 to 36% in 10 minutesThis was followed by a washout by a 1/2 min increase to 64% ACN, which was kept for 4.5 min. Flow rate was kept at 250 nL/min. Re-equilibration was done in parallel with sample pickup and prior to loading with a minimum requirement of 0.5 μL of 0.1% FA buffer at a pressure of 600 bar.

The mass spectrometer was running in data-dependent acquisition mode with the spray voltage set to 2 kV, funnel RF level at 40, and heated capillary at 275 °C. Full MS resolutions were set to 60 000 at m/z 200 and full MS AGC target was 3E6 with an IT of 25 ms. Mass range was set to 375–1500. AGC target value for fragment spectra was set at 1E5, and intensity threshold was kept at 2E5. Isolation width was set at 0.8 m/z and a fixed first mass of 100 m/z was used. Normalized collision energy was set at 33%. Peptide match was set to off, and isotope exclusion was on.

Raw MS files were analyzed by MaxQuant software version 1.6.0.17 using the Andromeda search engine. Proteins were identified by searching the higher-energy collisional dissociation (HCD)–MS/MS peak lists against a target/decoy version of the human UniProt protein database (release April 2017) using default settings. Carbamidomethylation of cysteine was specified as fixed modification, and protein N-terminal acetylation, oxidation of methionine, pyro-glutamate formation from glutamine, and phosphorylation of serine, threonine, and tyrosine residues were considered as variable modifications. The “maximum peptide mass” was set to 7500 Da, and the “modified peptide minimum score” and “modified maximum peptide score” were set to 25. Everything else was set to default values. The mass spectrometry proteomics data have been deposited to the ProteomeXchange Consortium via the PRIDE partner repository. Data are available via ProteomeXchange with identifier PXD018218 with the following reviewer account details: Username: Password: 4HEEtg6X

### 4.2 Data normalisation and differential analysis

In the phosphoproteomics dataset, 19285 unique phosphosites were detected across 18 samples. Visual inspection of the raw data PCA first 2 components indicated two major batches of samples. Thus, each batch was first normalised using the VSN R package^51,52^. We removed p-sites that were detected in less than 4 samples, leaving 14243 unique p-site to analyse. Visual inspection of the PCA first two components of the normalised data revealed that the first batch of samples could itself be separated in 3 batches (4 batches across all samples). Thus, we used the removeBatchEffect function of LIMMA to remove the linear effect of the 4 batches. Differential analysis was performed using the standard sequence of lmFit, contrasts.fit and eBayes functions of LIMMA, with FDR correction.

For the transcriptomics data, counts were extracted from fast.q files using the RsubRead R package and GRCh37 (hg19) reference genome. Technical replicates were averaged, and genes with average counts under 50 across samples were excluded, leaving 15919 genes measured across 22 samples. In order to allow for logarithmic transformation, 0 count values were scaled up to 0.5 (similar to the voom function of LIMMA). Counts were then normalised using the VSN R package function and differential analysis was performed with LIMMA package, in the same way as the phosphoproteomics data.

For the metabolomics data, 107 metabolites were detected in 16 samples. Intensities were normalised using the VSN package. Differential analysis was done using limma in the same manner as for phosphoproteomics and transcriptomics. All data is available at https://github.com/saezlab/COSMOS.

### 4.3 Footprint based analysis

TF-target collection was obtained from DOROTHEA A,B and C interaction confidence levels through the Omnipath webservice using the URL “http://omnipathdb.org/interactions?datasets=tfregulons&tfregulons_levels=A,B,C&genesymbols=1&fields=sources,tfregulons_level” (version of 2020 Feb 05). For the enrichment analysis, the viper algorithm^13^ was used with the limma moderated t-value as gene level statistic^53^. The eset.filter parameter was set to FALSE. Only TFs with at least 25 measured transcripts were included.

Kinase-substrate collection was obtained using the default resource collection of Omnipath, with the URL “http://omnipathdb.org/ptms?fields=sources,references&genesymbols=1” (version of 2020 Feb 05). For the enrichment analysis, the viper algorithm was used with the limma limma moderated t-value as phosphosite level statistic. The eset.filter parameter was set to FALSE. Only TFs with at least 5 measured transcripts were included. All data is available at https://github.com/saezlab/COSMOS.

### 4.4 Meta PKN construction

In order to propose mechanistic hypotheses spanning through signaling, transcription and metabolic reaction networks, multiple types of interactions have to be combined together in a single network. Thus, we built a meta Prior Knowledge Network (PKN) from three online resources, to incorporate three main types of interactions. The three types of interactions are protein-protein interactions, metabolite-protein allosteric interactions and metabolite-protein interactions in the context of a metabolic reaction network. Protein-protein interaction were imported from omnipath with the URL http://omnipathdb.org/interactions?genesymbols=1 (version of 2019 Feb 05), and only signed directed interactions were included (is_stimulation or is_inhibition columns equal to 1). Metabolic-protein allosteric interactions were imported from the STITCH database (version of 2019 November 06), with combined confidence score >= 900 after exclusion of interactions relying mainly on text mining.

For metabolic-protein interactions in the context of metabolic reaction network, Recon3D was downloaded from https://www.vmh.life/#downloadview (version of 2019 Feb 19). Then, the gene rules (“AND” and “OR”) of the metabolic reaction network were used to associate reactants and products with the corresponding enzymes of each reaction. When multiple enzymes were associated with a reaction with an “AND” rule, they were combined together as a single entity representing an enzymatic complexe. Then, reactants were connected to corresponding enzymatic complexes or enzymes by writing them as rows of Simple Interaction Format (SIF) table of the following form : reactant;1;enzyme. In a similar manner, products were connected to corresponding enzymatic complexes or enzymes by writing them as rows of Simple Interaction Format (SIF) table of the following form : enzyme;1;product. Thus, each row of the SIF table represents either an activation of the enzyme by the reactant (i.e. the necessity of the presence of the reactant for the enzyme to catalyse it’s reaction) or an activation of the product by an enzyme (e.i. the product presence is dependent on the activity of its corresponding enzyme). Most metabolite-protein interactions in metabolic reaction networks are not exclusive, thus measures have to be taken in order to preserve the coherence of the reaction network when converted to the SIF format. First, metabolites that are identified as “Coenzymes” in the Medical Subject Heading Classification (as referenced in the Pubchem online database) were excluded. Then, we looked at the number of connections of each metabolite and searched the minimum interaction number threshold that would avoid excluding main central carbon metabolites. Glutamic acid has 338 interactions in our Recon3D SIF network and is the most connected central carbon metabolite, thus any metabolites that had more than 338 interactions was excluded. An extensive list of Recon3D metabolites (pubchem CID) with their corresponding number of connections is available in supplementary table 2. Metabolic enzymes catalyzing multiple reactions were uniquely identified for each reaction to avoid cross-links between reactants and products of different reactions. Finally, exchange reactions were further uniquely identified according to the relevant exchanged metabolites, as to avoid confusion between transformation of metabolites and simply exchanging them between compartments. Finally, each network (protein-protein, allosteric metabolite-protein and reaction network metabolite-protein) was combined into a single SIF table. This network is available at http://metapkn.omnipathdb.org/.

### 4.5 Meta PKN contextualisation

Given a set of nodes with corresponding activities (−1, 0 or 1) and a causal PKN, CARNIVAL finds the smallest coherent signed subnetworks connecting as many of the given nodes as possible. CARNIVAL needs a set of starting and end nodes to look for paths in between. TFs, kinases and phosphatases absolute normalised enrichment scores greater than 1.7 standard deviation were considered deregulated. Coherently, metabolites with uncorrected p-values smaller than 0.05 were considered deregulated. These values were chosen as they allow to generate a set of input of comfortable size to run CARNIVAL. Then, we first set the deregulated kinases, phosphatases and TFs as starting points and deregulated metabolites as end points (forward run). This direction represents regulations first going through the signaling and transcriptional part of the cellular network and stops at deregulated metabolites in the metabolic reaction network. However, since metabolite concentration can also influence the activity of kinases and TFs through allosteric regulations, we also ran CARNIVAL by setting deregulated metabolites as starting points and deregulated TFs, kinases and phosphatases as end points (backward run). For the forward run, after 7200 second of run time, CARNIVAL yielded a network of 76 edges, a feasible solution which proved to be within the 11.24% gap from the optimal. For the backward run, CARNIVAL found a solution within the 2.44% gap from the optimal after 7200 second of run time, yielding a network of 177 edges.

Since there were no incoherences in the predicted activity signs between the common part of the two resulting networks, they were simply merged together, resulting in a combined network of 250 unique edges.

### 4.6 Coherence between CARNIVAL mechanistic hypotheses and omics measurements

To assess the robustness of CARNIVAL predictions, we used two different methods. First, the CARNIVAL network contains cases where a protein activity is modelled by CARNIVAL as up- or down-regulated under the control of a TF. If such hypotheses are correct, then one would expect to see the abundance of the corresponding transcript of the proteins to be coherently up or down-regulated (since the control of the TF is carried through regulation of transcript abundance) (Supplementary Figure 3). Thus, a True Positive (TP) is defined as a carnival node that is directly downstream of a TF and has the same sign (−1 or 1) as a significantly deregulated corresponding transcript. Transcripts with LIMMA moderated absolute t-values ranging between 0 and 2 were considered as significantly deregulated. Since CARNIVAL predictions are discrete (−1, 0, 1), we can’t make a classic receiving operator curve. Furthermore, we are lacking knowledge of True Negative. Indeed, node activities set to 0 by CARNIVAL cannot be interpreted because measurements and activities inputs only cover a fraction of the PKN and consequently most of the PKN nodes will be set to 0 by default. We showed that the TPR were relatively stable between 0 and 1.7, and more volatile between 1.7 and 2, likely due to the number of significantly deregulated transcripts considered becoming too small (see Supplementary Figure 4A). To estimate the baseline TPR of a random algorithm, we consider the following question : If i take any transcript that was measured and randomly assign it a value of 1 (or −1), what is the probability that the transcript will indeed be significantly up-regulated (or down-regulated), for given a t-value threshold. This probability can be simply estimated from the actual proportion of transcripts that are significantly up-related. For absolute t-values ranging between 0 and 1.7 (number of transcripts = 13), carnival TPR was consistently higher than the random baseline, but performed equal or worse than random above 1.7, again likely due to the number of significantly deregulated transcripts considered becoming too small.

Second, when multiple nodes are co-regulated by a common parent node in the CARNIVAL network, we can assume that the activity of the co-regulated nodes should be correlated. Thus, we create a correlation network with the TF and kinase/phosphatase activities estimated at a single sample level. To estimate the single sample level activities, normalised RNA counts and phosphosite intensities were scaled (minus mean over standard deviation) across samples. Thus, the value of each gene and phosphosite is now a z-score relative to an empirical distribution generated from the measurements across all samples. We used these z-scores as input for the viper algorithm to estimate kinase/phosphatases and TF activities at single sample level. Thus, the resulting activity scores in a sample are relative to all the other samples. Then, a correlation network was built using only tumor samples. Thus, the correlation calculated this way represents co-regulations that are supported by the available data in tumor (number of coregulations = 157. We defined the ground truth for co-regulations as over a range of absolute correlation coefficients between 0 and 1 with a 0.01 step. Thus, a True Positive here is a co-regulation predicted from the topology of the carnival network that also has a corresponding absolute correlation coefficient in tumor samples above the given threshold. Since defining a ground truth in such a manner can yield many false positives (a correlation can often be spurious), the TPR of COSMOS was always compared to a random baseline.

## 5. Acknowledgements

A.D. and E.G. were supported by the European Union’s Horizon 2020 research and innovation program (675585 Marie-Curie ITN ‘‘SymBioSys’’) to J.S.R., and were a Marie-Curie Early Stage Researcher. This work was supported by the Medical Research Council (MC_UU_12022/6 to C.F.). The Novo Nordisk Foundation Center for Protein Research is supported by Novo Nordisk Foundation grant number NNF14CC0001. J.V.O. was funded by a grant from Danish Council for Independent Research (8020-00100B) to partly support K.B.E. who was also supported in part by the Lundbeck Foundation (R193-2015-243). R.K. was supported by grants of the German Research Foundation (DFG: SFBTRR57, P30; SFBTRR219 C05, CRU344, P1), by a Grant of the European Research Council (ERC-StG 677448), a Grant of the State of North Rhine-Westphalia (Return to NRW), the BMBF eMed Consortia Fibromap, the ERA-CVD Consortia MEND-AGE, the Else Kroener Fresenius Foundation (EKFS) and the Interdisciplinary Centre for Clinical Research (IZKF) within the faculty of Medicine at the RWTH Aachen University (O3-11). C.K. was supported by the German Society of Internal Medicine (DGIM). Thanks to Hyojin Kim for her contribution to the original COSMOS logo design. Thanks to Denes Turei for his help with putting the metaPKN online.

## 6. Authors contributions

A.D. and J.S.R. designed the method. A.D. coded the pipeline and ran the analysis and interpreted results. E.G. developed and adapted CARNIVAL to the pipeline. K.B.E., D.B.B. and J.V.O. generated the phosphoproteomics dataset. E.M.J.B. performed final RNA-sequencing on Novaseq6000 platform. M.S. and A.S.H.C. performed the liquid chromatography mass spectrometry-based metabolomics analyses and processed the data. C.F., R.K. and J.S.R. supervised the project. A.D. wrote the manuscript with help from J.S.R.

## Supplementary Materials

**Supplementary Figure 1.**
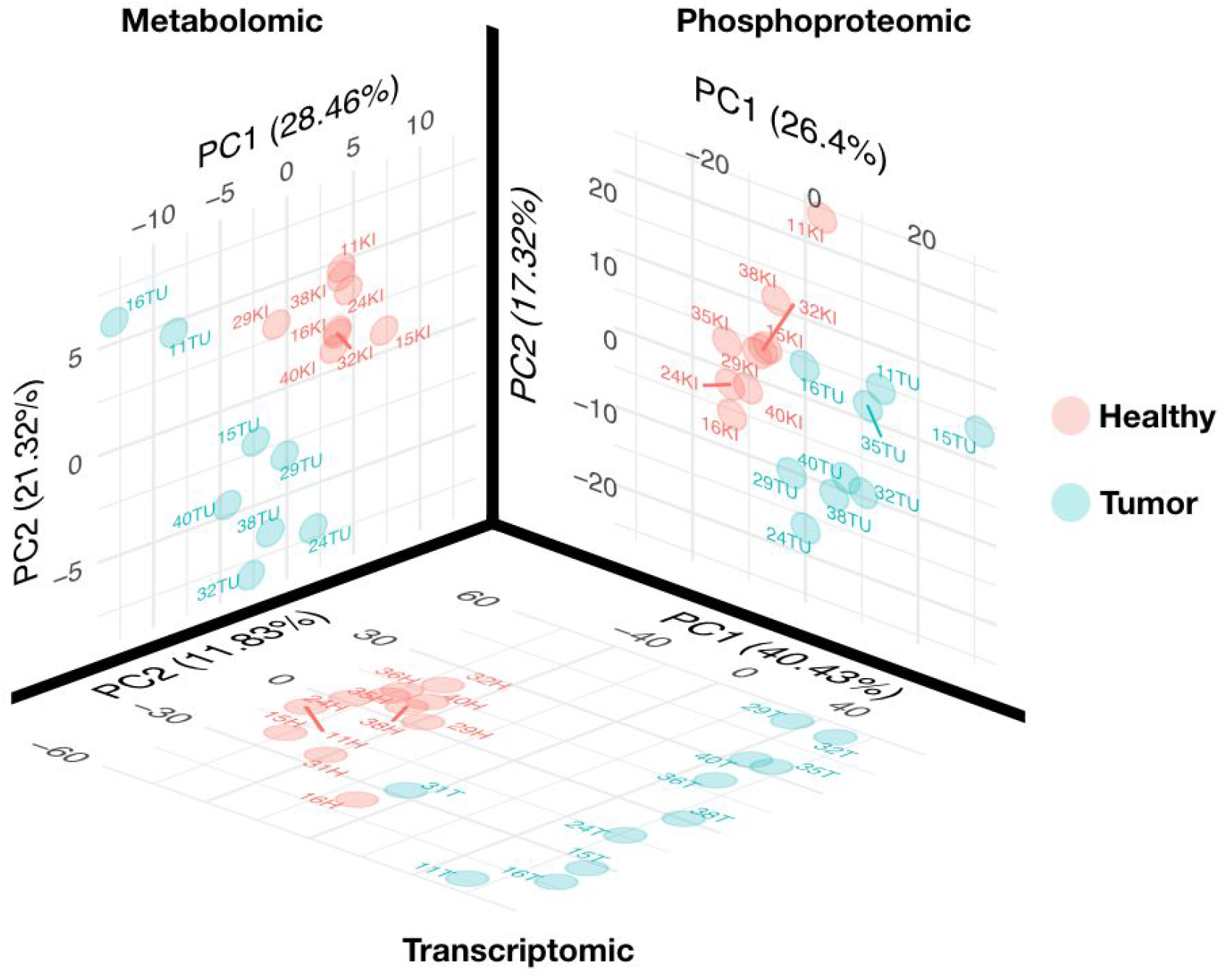
PCA of Metabolomics, Phosphoproteomics and transcriptomics datasets for tumor and healthy tissues samples. For each omics dataset, PCA is run independently on normalised datasets and the first two components are plotted. Each omics shows a clear separation between tumor and healthy tissue.

**Supplementary Figure 2.**
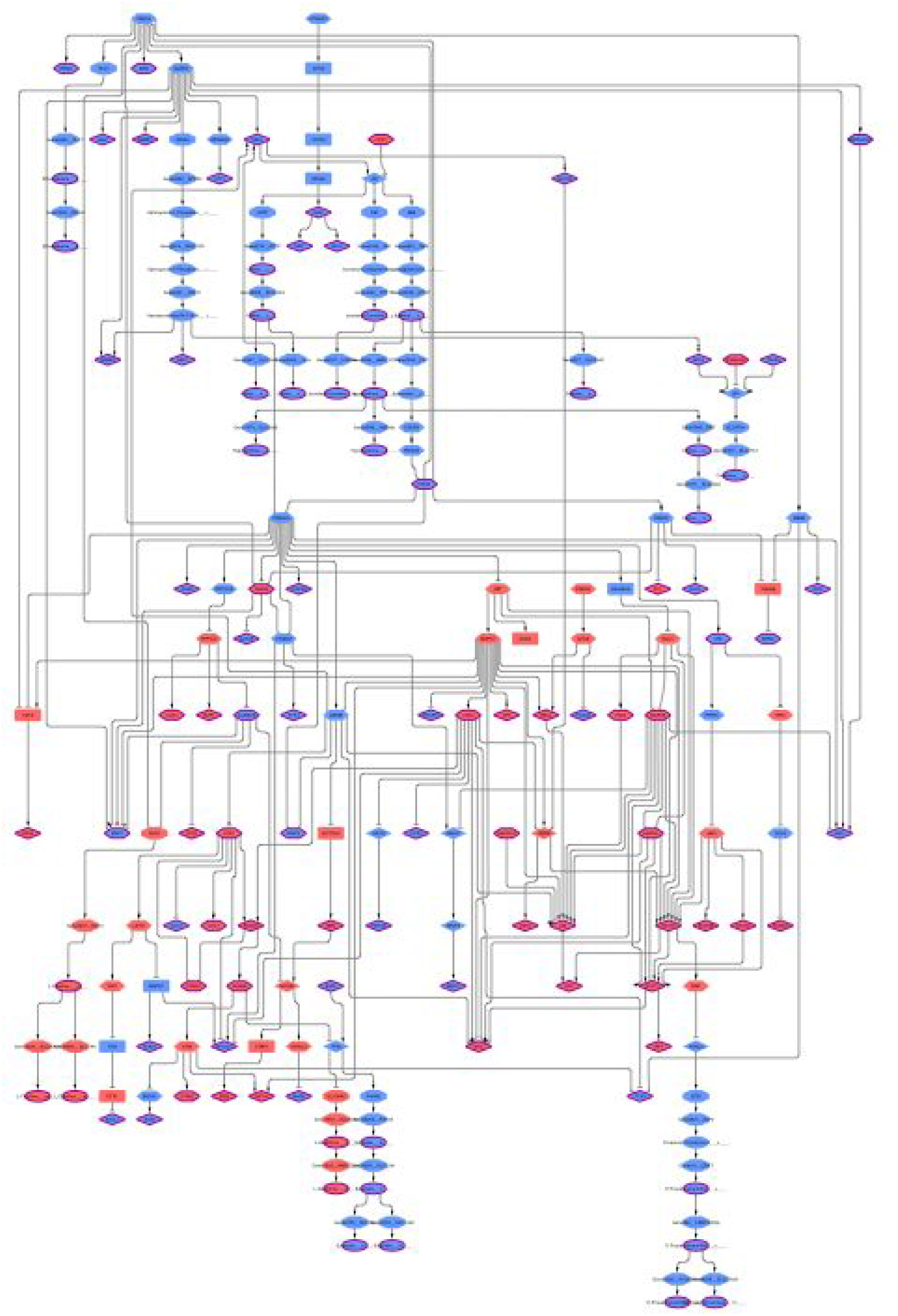
Causal network summarising the mechanistic hypotheses systematically generated by CARNIVAL. (see Figure 4 for legend). The network comprises 250 edges. It represents the propagation of signals connecting the deregulated kinases, phosphatases, TFs and metabolites in kidney cancer.

**Supplementary Figure 3.**
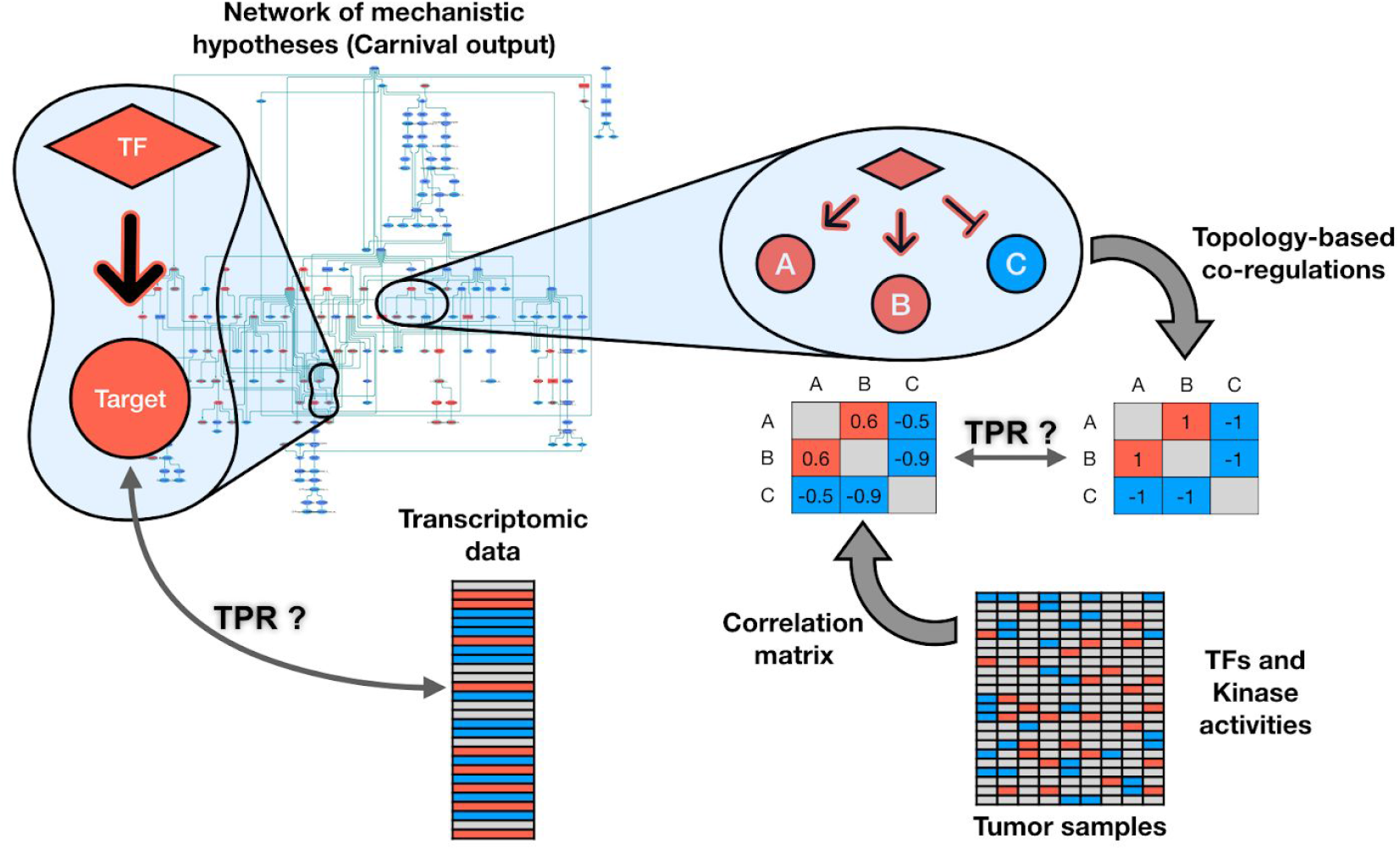
Coherence assessment between CARNIVAL hypotheses and underlying data. On the left, the predicted activity TF targets of the COSMOS network are compared to the actual t-value (tumor - healthy) of their corresponding transcript to determine true positive rate (TPR). On the right, coregulations predicted by COSMOS are compared against a correlation network of kinase/TF activities to determine TPR.

**Supplementary Figure 4.**
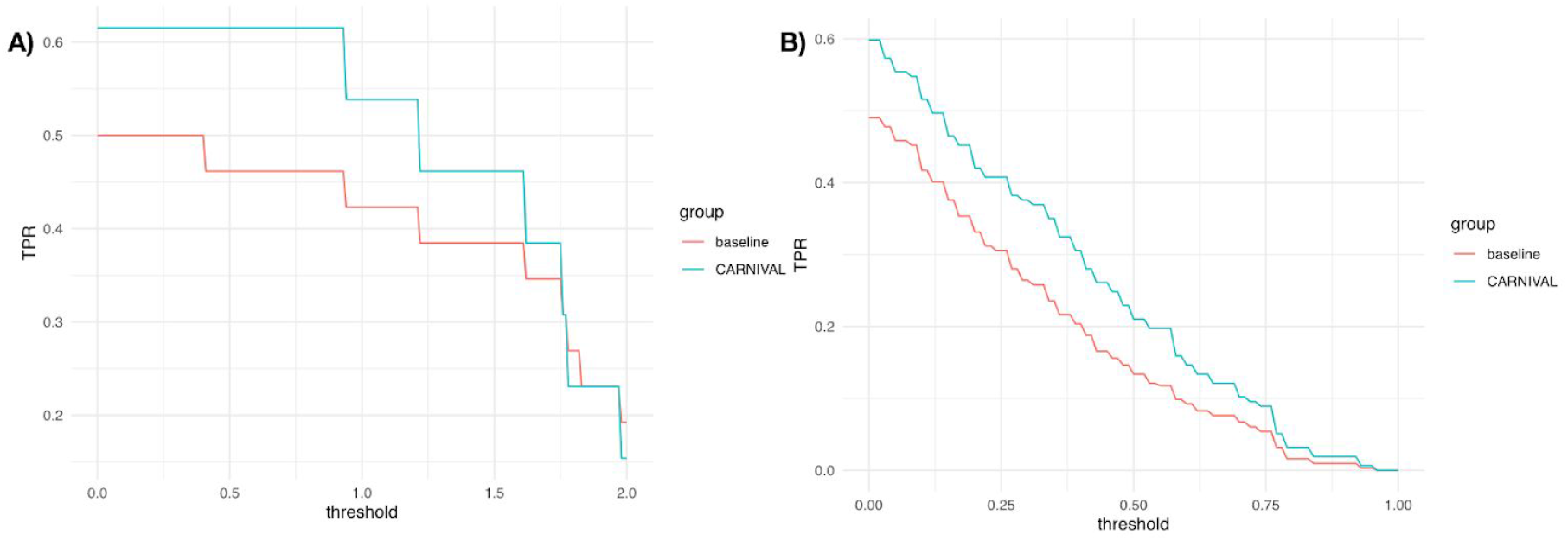
Exploration of TPR stability in function of the chosen t-value/correlation threshold. A) For TF/transcriptomics data coherence. True positive rates are estimated over a range of t-value between 0 and 2 with 0.1 steps. B) For the correlation/topology coherence. True positive rates are estimated over a range of Pearson correlation between 0 and 1 with 0.01 steps. In A) and B) COSMOS (CARNIVAL) performance is compared to a random baseline. COSMOS consistently outperforms the random baseline.

